# V1 receptive field structure contributes to neuronal response latency

**DOI:** 10.1101/2021.12.30.474591

**Authors:** Amin Vafaei, Milad Mohammadi, Alireza Khadir, Erfan Zabeh, Faraz YazdaniBanafsheDaragh, Mehran Khorasani, Reza Lashgari

## Abstract

The timing of neuronal responses is considered to be important for information transferring and communication across individual neurons. However, the sources of variabilities in the timing of neuronal responses are not well understood and sometimes over-interpreted. A systematic variability in the response latencies of the primary visual cortex has been reported in presence of drifting grating stimulus. Whereas the response latencies are systematically dependent on stimulus orientation. To understand the underlying mechanism of these systematic latencies, we recorded the neuronal response of the cat visual cortex, area 17, and simulated the response latency of V1 neurons, with two geometric models. We showed that outputs of these two models significantly predict the response latencies of the electrophysiology recording during orientation tasks. The periodic patterns created in the raster plots were dependent on the relative position of the stimulus rotation center and the receptive-field sub-regions. We argue the position of stimulus is contributing to systematic response latencies, dependent on drifting orientation. Therefore, we provide a toolbox based on our geometrical model for determining the exact location of RF sub-regions. Our result indicates that a major source of neuronal variability is the lack of fine-tuning in the task parameters. Considering the simplicity of the orientation selectivity task, we argue fine-tuning of stimulus properties is crucial for deduction of neural variability in higher-order cortical areas and understanding their neural dynamics.

## Introduction

The primary visual cortex (V1) contains non-orientation selective (NOS) and orientation-selective neurons (OS). NOS neurons have a center-surround receptive-field that is ON or OFF center. OS neurons have at least two antagonist regions that respond either to dark (OFF) or light (ON) stimuli ^1^. Finding the exact location of the receptive field (RF) and its sub-regions is essential for many system and cognitive experiments, especially those that deal with drifting stimuli or neuronal spike latency.

The neuronal spike latency of V1 neurons is subject to variability similar to other neuronal activity in cortex ^2,3^, however, it is not known how much of this temporal variability is stimulus driven^4^ or how much of it is due to ongoing oscillation ^5,6^ or bottom up influences ^7,8^. A series of studies propose there is a functional relationship between the latency of V1 response and the eccentricity from preferred properties of the neuron, suggesting stimulus properties play a role in the generation of response latency properties. Accordingly, identifying the optimal stimulus is beneficial to deduce neuronal response variability and reveal the actual neural dynamics of the target neuronal population.

For a long time, researchers have been using Hartley ^9^, sparse noise ^10^, m-sequence ^11^, or two-dimensional Hermite ^12^ stimuli to map retinotopic RFs (segregated into ON and OFF regions) using reverse correlation. However, because of spontaneous activity and variability in neural responses, there is always an error in the estimation of RF’s sub-region locations. It also has been noted that many of these methods are not reliable for estimating RF, as different mapping methods lead to different results ^12,13^. Here we propose a method for determining the exact location of RF regions in V1 neurons (both simple and complex) that is not dependent on individual spikes and is instead based on the latency patterns in the raster plot.

We have seen diverse but consistent patterns of latency in the raster plots of V1 neurons responding to drifting grating stimuli (fig. 1c/d). Figure 1e illustrates the distribution of half bandwidth for these 63 OS neurons. The median of the distribution is 26 degrees, which is consistent with earlier studies ^14–16^. These patterns have been previously observed by other researchers as well ^17–19^, although there is not an agreed-upon explanation of the underlying mechanisms for them. Shriki et al^18^ suggest that V1 neurons use these latency patterns to encode the orientation of the stimulus. We use a geometric model to show that, although there is a strong correlation between orientation and latency patterns (fig. 1f/g), it is only an artifact of the already-known functionality of neural activity in V1 cells. We show that the latency of neural response to drifting grating stimuli depends on the orientation of the stimulus and the relative location of the stimulus rotation center into the RF. Consequently, developed a method to find the accurate location of the RF sub-regions, based on these latency patterns, with an error of almost 1/3 of Hartley mapping. Using single and multi-electrode recordings from cat and monkey V1, we show that our model’s output is consistent with the neural activity and can be used in action for accurate RF mapping.

**Figure 1.**
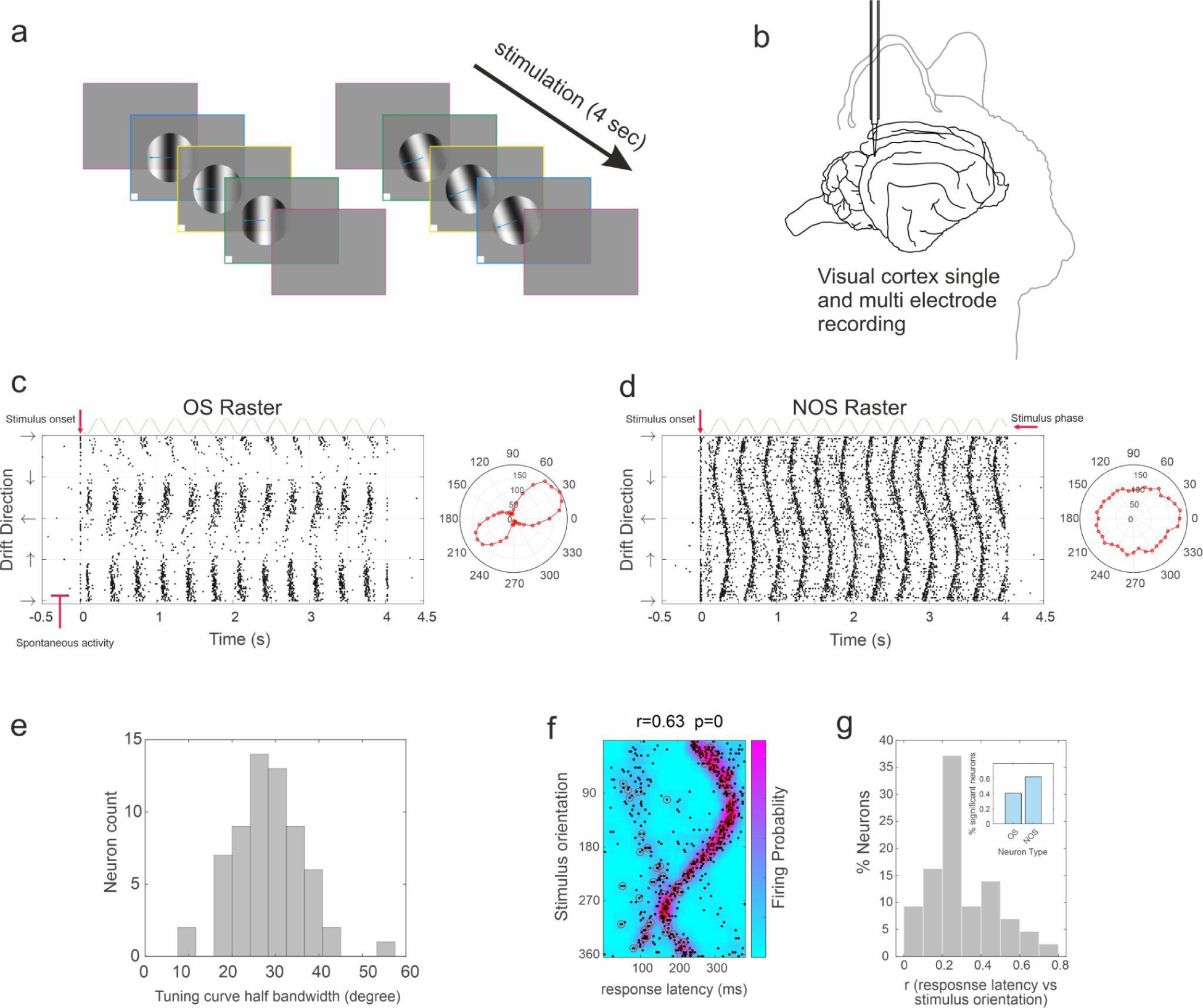
Electrophysiological recordings and systematic latency dependence to stimulus orientation. (a) Moving Drifting grating stimulus with the spatial frequency of (0.1 c/degree) and (3 Hz) temporal frequency. The drift direction changed in each trial (4s). (b) FHC tungsten single-electrodes were vertically introduced through the cat visual cortex (area 17) and receptive fields were measured at each cortical site with Hartley stimuli. (c/d left) Spike raster of two representative neurons, (c) orientation selective and (d) non-orientation selective. Horizontal and vertical axes represent the timing of the task (respect to stimulus onset) and drifting direction. The periodic characteristic of firing rate is due to changes of stimulus phase. (c/d right) the tuning curve of the representative neurons in response to different stimulus orientations. The turning curve of the orientation selective neuron indicates significant selectivity for 30 degree and 210-degree stimuli (p<0.05, ryleigh test). (e) Distribution of half bandwidth for 63 OS neurons. The median of distribution is 26 degrees, which is consistent with the earlier reports. (f)The firing probability of the representative neurons indicates a circular linear correlation between response latency and stimulus orientation (g) Distribution of circular-linear correlation coefficient (r) between stimulus orientation and response latency. Majority of neural response latencies are significantly correlated with stimulus orientation. (Upper right corner represents the percentage of significant selective neurons) The stimulus dependent response latency is more ubiquitous in NOS neurons.

There is a widespread belief about orientation selectivity which is “The orientation selectivity of simple cells derives directly from the shape of their receptive field (Carandini et al^20^, Fig. 5): ON and OFF sub-regions are elongated, so their preferred stimulus is similarly elongated” and the preferred orientation could be predicted from RF structure ^20–24^.

Our model works based on sensitivity to a single point on the screen for NOS neurons and sensitivity to two points for OS neurons. The precise operation of this model shows that V1 neurons require at least two spatially offset circular sub-regions for orientation selectivity, which must be activated simultaneously. Indeed, the mechanism of orientation selectivity in V1 neurons depends on the existence of two opposing circular sub-regions (ON and OFF) and not the ellipsoid form of their sub-regions. Also, we suggest that the reason behind the ellipsoid form of sub-regions calculated by some RF mapping tasks is because of the nature of RF mapping tasks and not the real shape of sub-regions. Consider that there are different RF maps as a result of using different RF mapping tasks. For example, there are no elongated (ellipsoid-shaped) sub-regions using sparse noise RF mapping tasks with only white or black stimuli ^13,17,25^. Our finding is consistent with the fundamental law of geometry that at least two points are needed to draw a line as V1 neurons need at least two circular sub-regions for orientation selectivity and detecting edges.

## RESULTS

### Geometrical Model for NOS neurons

Here we basically explain the functioning of the model based on NOS neurons. NOS neurons have center-surround RFs, so to demonstrate our model we simulate this type of neuron based on sensitivity to the single point area considering the receptive field center on the screen. In (fig. 2c/d) we have shown inaccurate positioning of the grating stimulus rotation center (Yellow Dot A) relative to the RF center (hatched red circle) in two different orientations. Figure 2a illustrates the center of ON-center RF by a hatched red circle, in which crossing the grating white line from RF center (point B) will activate the neuron.

**Figure 2.**
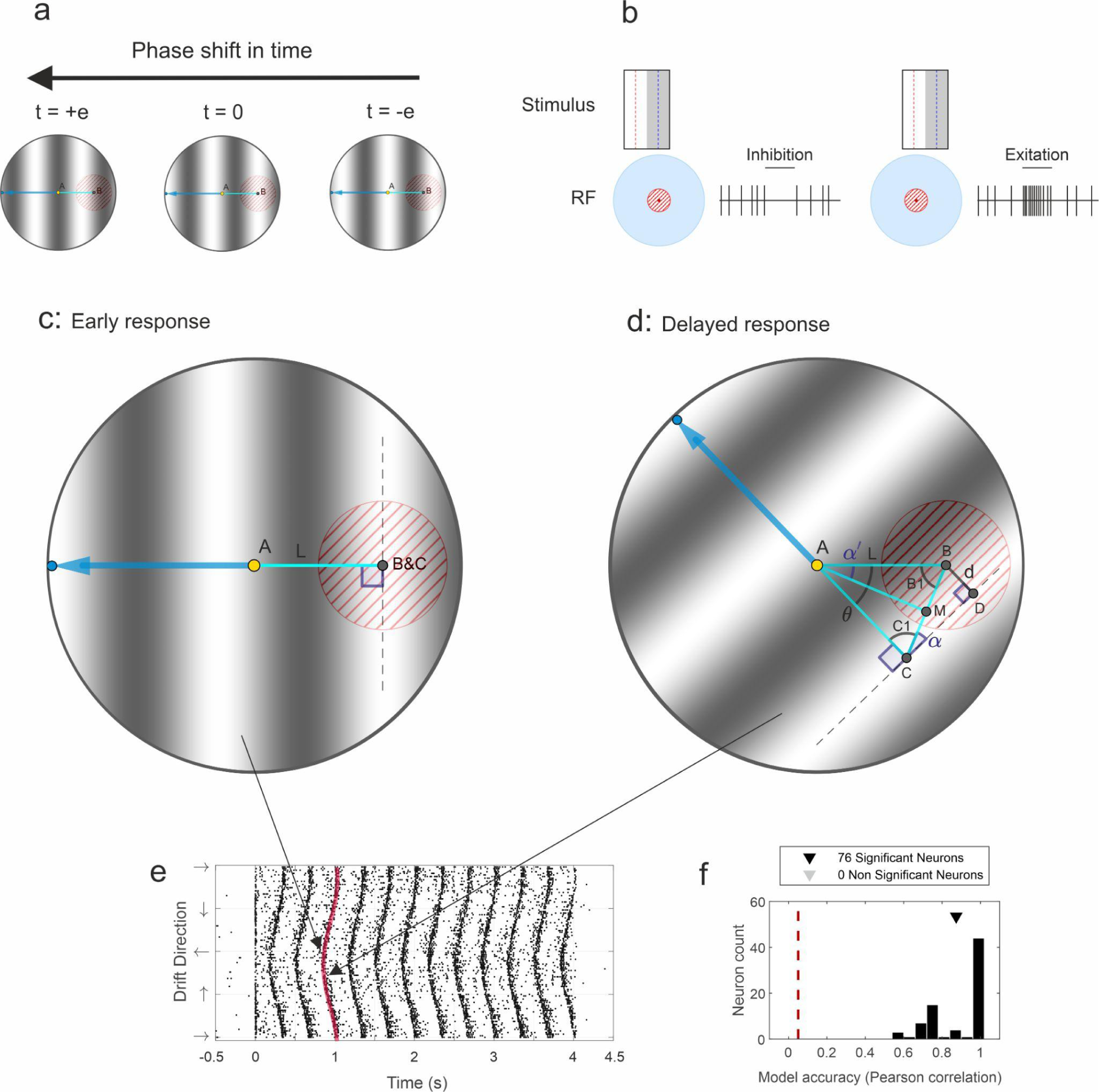
Geometric model representation of path difference as a function of drift direction. Our model hypothesizes the mismatching between stimulus center and RF is contributing to systematic response latency, dependent on drifting orientation. (a) Illustration of stimulus drifting (phase development with time). From left to right in each snapshot, the RF center (illustrated with a dashed red circle) experiences different stimulus polarities which end up with different firing of the neuron. (b) Illustrating the maximum and minimum firing rate of neurons in response to different stimulus phases. (c) Model for NOS neurons that shows RF center and drifting stimulus at initial phase (time t=0). Here the stimulus rotation center (yellow point A) is positioned inaccurately left of the RF center. By crossing the stimulus white line from the RF center, the neuron is activated at time t=0. The blue arrow shows the drifting direction, the dashed line indicates the grating white line center, and line L indicates the distance between the stimulus rotation center and RF center. (d) In another drift direction at time t=0, to activate the neuron, you need to wait until the stimulus drifts a distance *d*. This introduces latency in neuronal activation as a function of stimulus drift direction. (e) Creation of latency patterns in the raster plot. We have shown the response latency for two different drift directions on the raster plot. (f) Distribution of Pearson correlation between sinusoidal model fit and actual response latency of individual neurons. The distribution is calculated over all recorded NOS neurons. The red vertical line represents the significance threshold.

Consider that at time t=0, the stimulus is positioned such that the neuron is activated (positioning of the grating white line on the RF center, fig. 2a/c). If we repeat the experiment but with a rotated stimulus (fig. 2d), the neuron no longer activates at t=0. Rotation of grating stimulus around point A (stimulus center) creates (*d*) displacement of grating white line relative to the RF center. Instead, for neuronal activation, we need to wait until the stimulus drifts a distance (*d*). This geometric displacement creates different response latency in different orientations (fig. 2e). The concept and driving of geometrical equations is described based on table 1 (in methods) and parameters shown in (fig. 2d). The grating displacement (*d*) relative to the rotation angle is measured and described below:

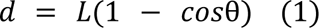

**Table 1:** Driving of geometrical equations.

|  |  |  |
| --- | --- | --- |
| $AB = AC \rightarrow C_1 = B_1$ | $ABC$ is the equilateral triangle | (4) |
| $CM = BM$ | $M$ is the middle point of $BC$ | (5) |
| $\alpha = \frac{\theta}{2}$ | $AM$ is the bisector of $(\theta)$ | (6) |
| $C_1 = \frac{\pi}{2} - \alpha' = \frac{\pi - \theta}{2}$ | Sum of angles of a triangle ($ABC$, $CBD$) equals $\pi$ | (7) |
| $BM = L \times \sin[\frac{\theta}{2}]$ | Based on trigonometric equations | (8) |
| $CB = 2L \times \sin[\frac{\theta}{2}]$ | $CB$ is twice as much as $BM$ | (9) |
| $d = CB \times \sin(\alpha') = 2L \sin^2[\frac{\theta}{2}] = L(1 - \cos\theta)$ | Based on trigonometric equations | (10) |

Where *L* is the distance between stimulus center (point A) and RF center (point B), θ is the rotation angle (i.e., orientation of the stimulus) and *V* is drift velocity. Therefore, spike latency would be:

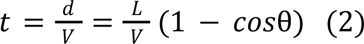

Also, the amplitude of the sinusoidal pattern would be:

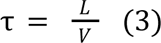

This predicts that, if using drifting grating stimuli and change the orientation gradually in each trial, the patterns created in the raster plots of NOS V1 neurons (ON or OFF-center neurons) will follow a sinusoidal function of the orientation, a pattern that has been observed before ^17,18^. The accuracy of this prediction is shown in (fig. 2f) and indicates that the actual latency patterns in the raster plot of NOS neurons are sinusoidal. This sinusoidal relationship disappears if the receptive field is exactly located on the center of the stimulus, where *L* = 0 and the latency pattern does not change with the orientation. Also, figure 1 in supplementary shows increase in response latency amplitude as a result of increasing the distance between stimulus rotation center and RF center. Therefore, we can use this fact to exactly find the RF location for non-orientation selective neurons.

### Application of method on NOS neurons

Here the neuronal response latency has shown in two conditions (fig. 3b/c), based on our model. The drifting grating stimulus has shown in (fig. 3a left) at the start of the task (time = 0). Red and blue dotted lines show the center of stimulus white and black lines. Also (fig. 3a right) shows the center-surround ON center RF. By positioning the stimulus rotation center on the RF center (fig. 3b), we have a constant distance between the stimulus white line (red dotted line) to the RF center in different drift directions. Indeed, neuronal response latency would be equal in all drift directions. This result also could be obtained by [equation.2] and the value of zero for L. In (fig. 3b) can see the neuronal response latency for inaccurate positioning of the stimulus rotation center. In this positioning, different drift directions make different distances between the stimulus white line (red dotted line) to the RF center. So, we have different neuronal response latency in different drift directions that this latency could be obtained by [equation.2] and the non-zero value of L.

**Figure 3:**
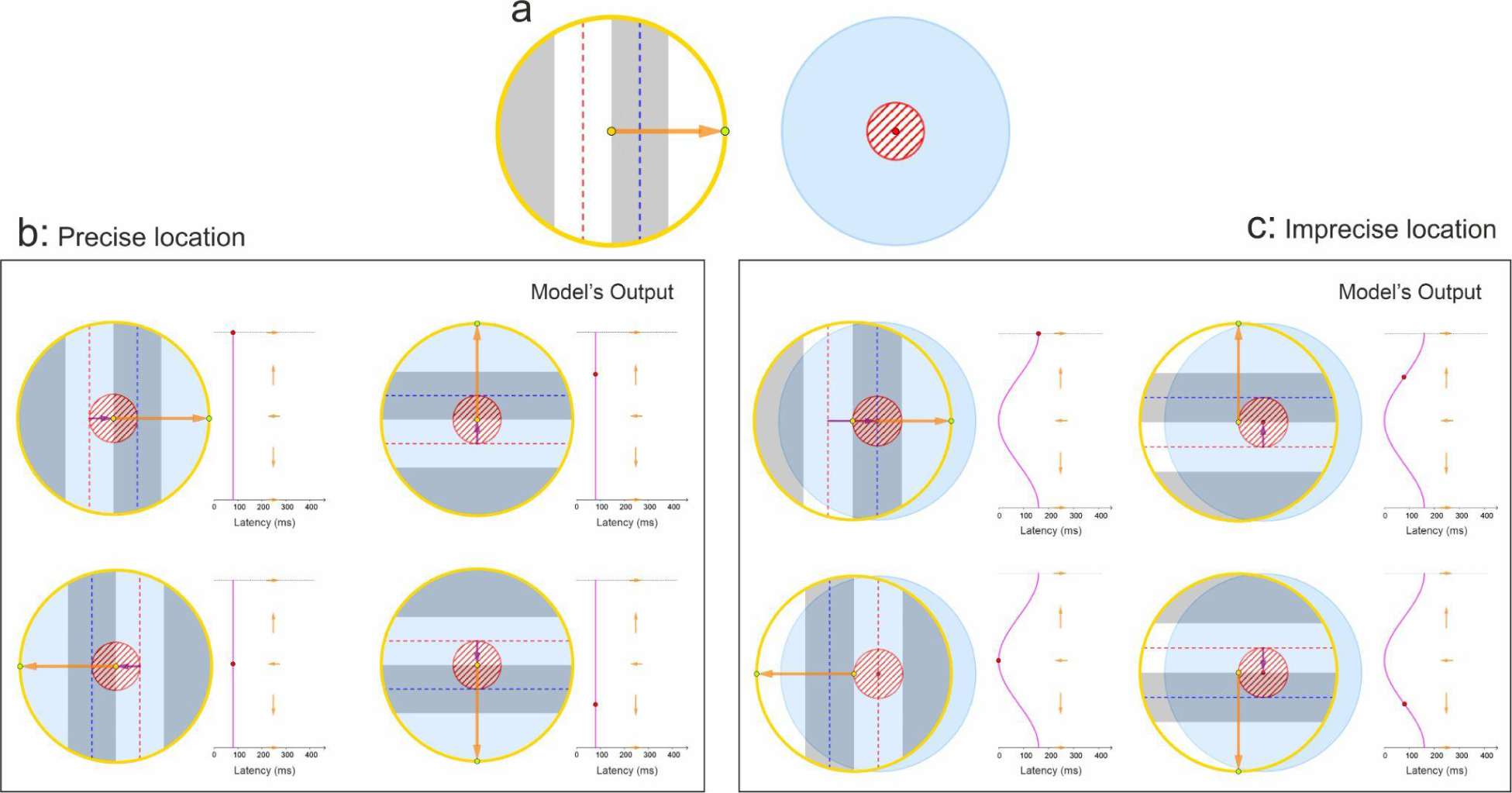
Modeling of response latency for NOS neurons. (a) Cartoon illustration of grating stimulus (left) and NOS ON-center RF (right). Red and blue dotted lines (left) indicate the center of white and black lines in the grating stimulus. The orange arrow shows the stimulus drift direction. (b) Cartoon positioning of stimulus on the RF (left) and created response latency based on model (right). By positioning the stimulus rotation center (yellow dot) exactly on the RF center (center of the red hatched circle), we have constant response latency in all drift directions (red dot on vertical purple line). For neuronal activation, the stimulus should drift distance from the stimulus white line center to the RF center (purple arrow). Here we have shown latency in four different drift directions. (c) Inaccurate positioning of the stimulus relative to the RF center creates the sinusoidal pattern in the raster plot. Here we have shown the creation of sinusoidal patterns in four different drift directions as a result of the geometric model.

To evaluate the accuracy of the model, we compared the predicted results of the model with the actual data recorded from Cat and Monkey V1 neurons, responding to moving bars and the drifting grating task. In recordings of non-orientation selective V1 neurons in the cat experiment, we manually determined the approximate location of the receptive field and presented optimum drifting grating stimuli (orientation task) in the estimated location (fig. 4a right green dot), which creates sinusoidal patterns in the neurons raster plot (fig. 4a left). According to the prediction of the model, the sinusoidal pattern in the raster-plot indicates the inaccuracy of the manually estimated receptive field location. To identify the exact location of the receptive field for the neurons with sinusoidal patterns in their raster plots, we first fitted the equation [equation. 2] on the raster plot to find L (fig. 4b). L is the correction displacement distance and after determining L we need the displacement direction that is the direction of the drifting with the highest response latency. This can be done based on our written application or manually. So, we can find the exact location of RF that is shown by the yellow dot in (fig. 4a/c milled and right). As shown, the yellow dot completely matches the RF center (fig. 4a middle) calculated by reverse correlation (explained in the methods section) in which the red area shows the center of ON-center RF. Green dot in (fig. 4a/c middle and right) shows the stimulus rotation center.

**Figure 4:**
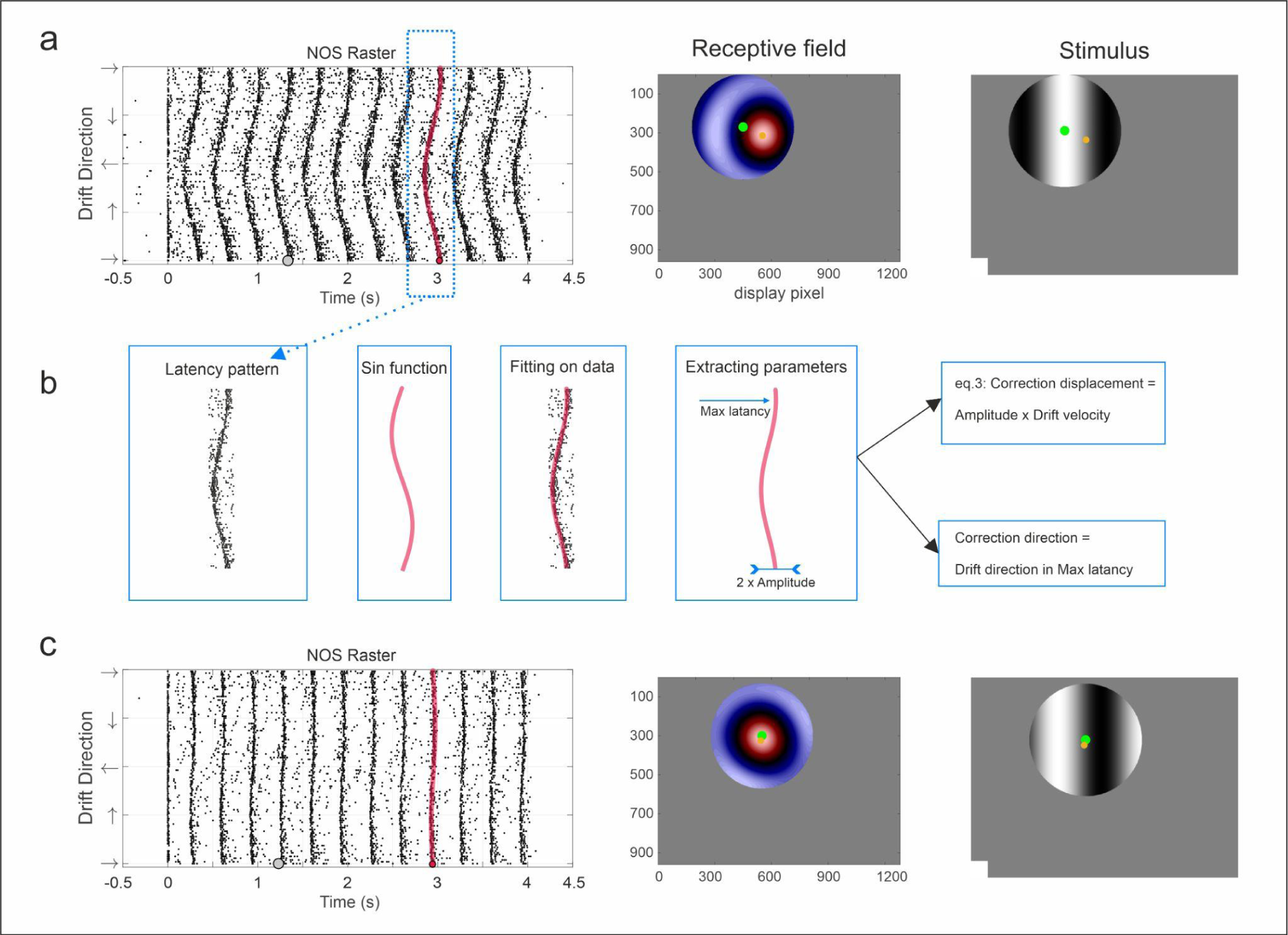
Evaluation of a geometric model for cat NOS V1 neurons. We used patterns in the raster plot to find the RF center accurately. (a) Illustration of a sample NOS raster plot (left), the neuron’s RF (middle), and the positioning of stimulus on the screen (right). We mapped the RF-based on reverse correlation (methods) and the red circular region in the map indicates the ON center NOS neuron. In the middle and right, green and yellow points show the stimulus rotation center and RF center. Indeed, the green point is determined manually, while the yellow point is obtained based on the latency pattern in the raster plot. The gray point on the raster plot (left) also shows the phase and the direction of the grating stimulus (right) at the appropriate time point in which the neuron starts to fire. (b) Illustration of applying the method to the real data. By extracting the pattern from the raster and fitting the sinusoidal function (based on the model’s formula), we can find the direction in which the neuron shows maximum latency and also amplitude of the pattern. Based on these parameters and the location of the green point, we can find the accurate location of RF (yellow point). (c) By positioning the stimulus rotation center (green point) on the RF center (yellow point), we can see vertical lines in the raster plot. It shows that we can accurately find the RF center based on the sinusoidal pattern in the raster plot.

So, we replaced the stimulus position in the exact RF center (yellow dot in fig. 4a/b milled and right), then we repeated the experiment to see if the sinusoidal pattern disappeared (fig. 4c left). This new accurate position creates vertical lines in the raster plot. As shown, the sinusoidal raster pattern depends completely on the relative location of RF and stimulus rotation center.

### Application of the method for OS neurons

To further provide evidence for the proposed model, we here used the model to explain the patterns observed in orientation-selective neurons. OS neurons have RFs with two controversial sub-regions, so we simulate these neurons based on sensitivity to two points on the screen. The drifting grating stimulus has shown in (fig. 5a left) at the start of the task (time = 0). Red and blue dotted lines show the center of white and black lines in the stimulus. Also (fig. 5a right) shows the two sub-regional RF for OS neurons. Consider that our model only works based on two circular sub-regions (fig. 5a right) and here we have also shown the dotted ellipsoid form of RF sub-regions for a better understanding of our model. We should note that this ellipsoid form of sub-regions only occurs because of the nature of RF mapping and plays no role in orientation selectivity and as we have shown, only circular sub-regions are enough for orientation selectivity. Following the same paradigm on non-orientation selective neurons, we show that each sub-region in the receptive field, if taken into account in isolation, activates with a latency that follows a sinusoidal pattern when presented with an optimum drifting grating stimulus. So, there are two different sinusoidal latency patterns for ON and OFF regions. In this model, an orientation-selective neuron activates when both sub-regions are activated at the same time, which means that we can see spikes in the regions of the raster plot where the sinusoidal patterns collide. In (fig. 5b/c) we have shown the neuronal response latency in two conditions, based on our model. Positioning of the stimulus rotation center relative to the RF sub-regions creates different patterns in the raster-plot. In (fig. 5b), we have shown patterns of latency for both subregions as a result of positioning the stimulus rotation center on the RF center (between two sub-regions). In this model, neuronal activation only occurs where the sinusoidal patterns collide. Also, we have shown different positioning of the stimulus rotation center relative to the RF (fig. 5c) and as a result, we can predict oblique activation zones (overlap of latency patterns) in the raster plot. Different types of patterns could occur in the raster-plot and it depends on the positioning of the stimulus rotation center. We have shown another type of pattern in supplementary (fig. 3).

To evaluate the accuracy of the model, we compared the predicted results of the model with the actual data recorded from Cat V1 neurons, responding to the drifting grating task. In recordings of orientation-selective V1 neurons in the cat experiment, we manually determined the approximate location of the receptive field (fig. 6a right) and presented optimum drifting grating stimuli (orientation task) in the estimated location, which creates oblique patterns in the neuron’s raster plot (fig. 6a left). By fitting our model’s output (fig. 6a left. blue and red lines) on the created pattern we can find the accurate location of RF sub-regions. Blue and red dots in (fig. 6a middle and right) show the accurate center of RF sub-regions as the model’s output. The green dot in (fig. 6.a/b middle and right) shows the stimulus rotation center. Again, for validating our model’s performance, we put the stimulus rotation center on the model’s output (fig. 6a/b middle and right yellow dot) and repeated the orientation task (fig. 6b). This displacement changes patterns in the raster plot and illustrates that the model also works accurately for OS neurons. Indeed, as our model accurately creates activation zones in raster-plots (overlap of latency patterns) only based on two circular sub-regions, we could claim that orientation selectivity could only occur based on two circular sub-regions and not the ellipsoid form of sub-regions.

**Figure 5:**
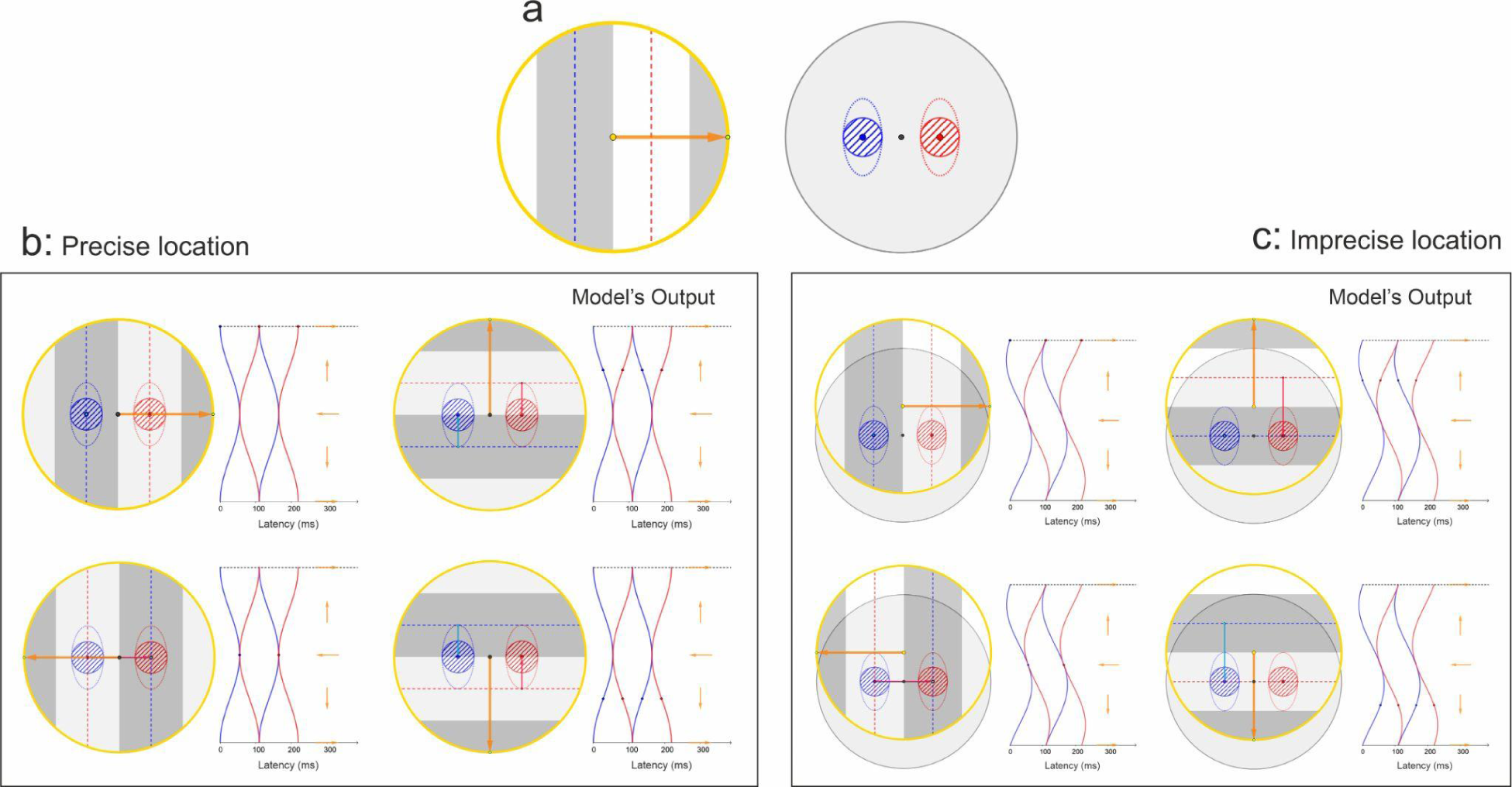
Modeling of response latency for OS neurons. (a) Cartoon illustration of grating stimulus (left) and OS RF (right). Red and blue dotted lines (left) indicate the center of white and black lines in the grating stimulus. The orange arrow shows the stimulus drift direction. Blue and red circular sub-regions show ON and OFF regions and dotted ellipses are only for better understanding. Note that the model only works based on two circular sub-regions. (b) Patterns are created by the positioning of the stimulus rotation center (yellow dot) exactly on the RF center. Created patterns in the raster plot depend on the positioning of the stimulus rotation center relative to the RF sub-regions. Like the NOS model, we have sinusoidal latency patterns for each sub-region that is shown by the blue and red lines, but activation occurs on the overlap of these patterns. (c) Patterns are created by the positioning of the stimulus rotation center (yellow dot) on an inaccurate position (top of RF center).

**Figure 6:**
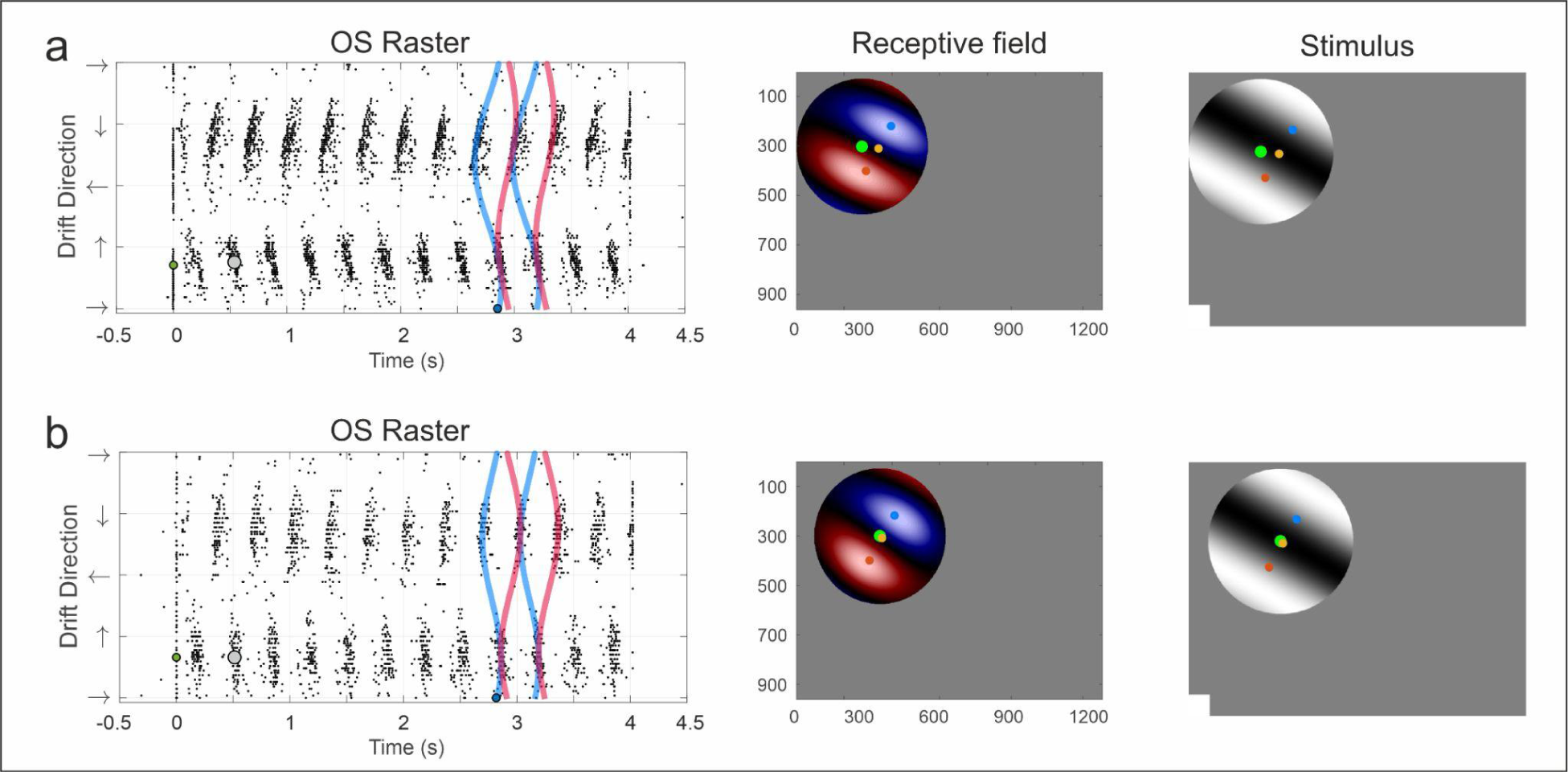
Evaluation of a geometric model for cat OS neurons. We used patterns in the raster plot to find the RF subregions accurately. (a) Illustration of a sample OS raster plot (left), the neuron’s RF (middle), and the positioning of stimulus on the screen (right). We mapped the RF-based on reverse correlation (methods) in which blue and red ellipsoids are OFF and ON sub-regions. In the middle and right, green and yellow points show the stimulus rotation center and RF center. Indeed, the green point is determined manually, while the yellow point is obtained based on the latency pattern in the raster plot. Also, the blue and red points show the accurate RF sub-regions, obtained by the model. The gray point on the raster plot (left) also shows the phase and the direction of the grating stimulus (shown in right) at the appropriate time point in which the neuron starts to fire. By fitting the model’s output (blue and red sinusoidal lines on raster plot) on the real neuronal pattern, we can find the exact location of RF sub-regions (blue and red dot middle and right) as a result of the model. (b) By positioning the stimulus rotation center on the RF center, we can see vertical patterns in the raster plot. It shows that we can accurately find the RF sub-regions based on the pattern in the raster plot. As illustrated on raster plots, we can see neuronal activation at the intersection of two sinusoidal patterns. As our model works based on two circular sub-regions, simultaneous activation of two circular ON and OFF subregions will activate the OS neurons.

### Hartley error correlation based on geometrical method

To study the accuracy of RF mapping generated with our method in comparison with the Hartley stimulus, for V1 NOS neurons, originated from monkey and cat, we first mapped the RF location using Hartley stimulus (fig. 7a), and then the optimum drifting grating stimulus was presented in the predicted location (fig. 7a blue dot). This protocol produced sinusoidal patterns in the raster plot (fig. 7b), which means that the location estimated by Hartley was not accurate. Using the geometric method, we were able to correct the estimation of the location of the receptive field (fig. 7a green dot), validated by raster plots with vertical lines rather than sinusoidal patterns (fig. 7c).

**Figure 7:**
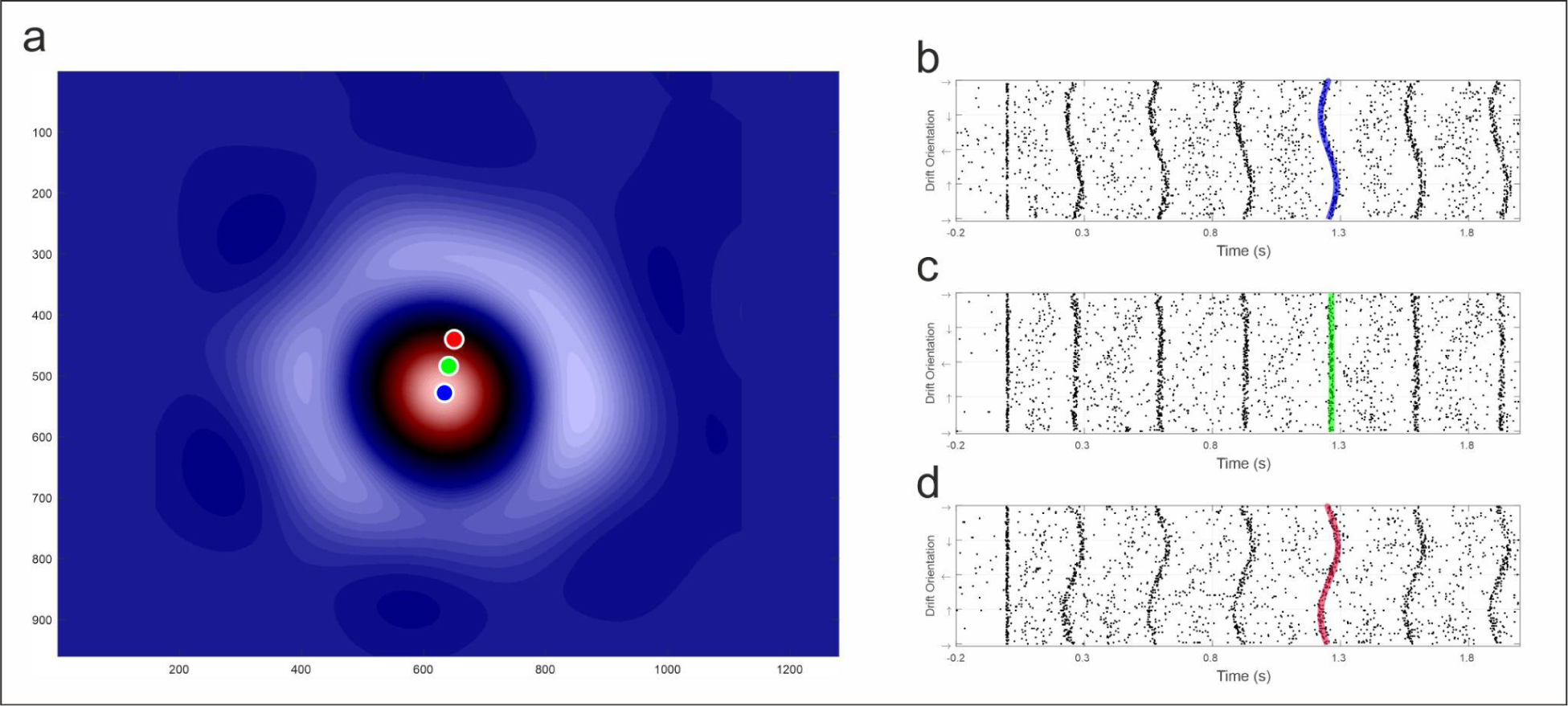
Error in Hartley RF mapping. (a) RF map for a NOS neuron calculated by Hartley mapping. The red circular region in the map indicates the ON center NOS neuron. Blue, green, and red dots on the map shows the drifting grating rotation center for part b, c, and d. (b) Positioning the drifting grating rotation center on the center of the Hartley map (blue point) creates a sinusoidal pattern in the raster plot. It shows that the center of Hartley is not an accurate estimation for the RF center. (c) Based on the sinusoidal pattern in b, we can find the accurate location of RF (green point). To test the accuracy of the model, we put the stimulus rotation center on the green point. So, we can see vertical lines in the raster plot as a result. It shows the accuracy of our method for RF mapping. (d) Reverse positioning of stimulus rotation center (red point) creates reverse sinusoidal patterns in the raster plot.

To quantify the accuracy of our method and compare against Hartley, we measured the amplitude of the sinusoidal pattern (τ) in the raster plot of V1 NOS neurons, when presenting stimuli in the location estimated by Hartley, as well as the locations corrected using our method. Based on the amplitude of the sinusoidal pattern, we calculated the spatial RF mapping error and normalized the error, dividing by the FR sizes for each neuron (fig. 8 dataset 1 & 2). We calculated error for 270 recorded neurons from monkeys and 11 neutrons from cats. Figure 8 shows that using our proposed method, the error in the estimated RF location can be decreased to 1/3 of that of the Hartley method.

**Figure 8:**
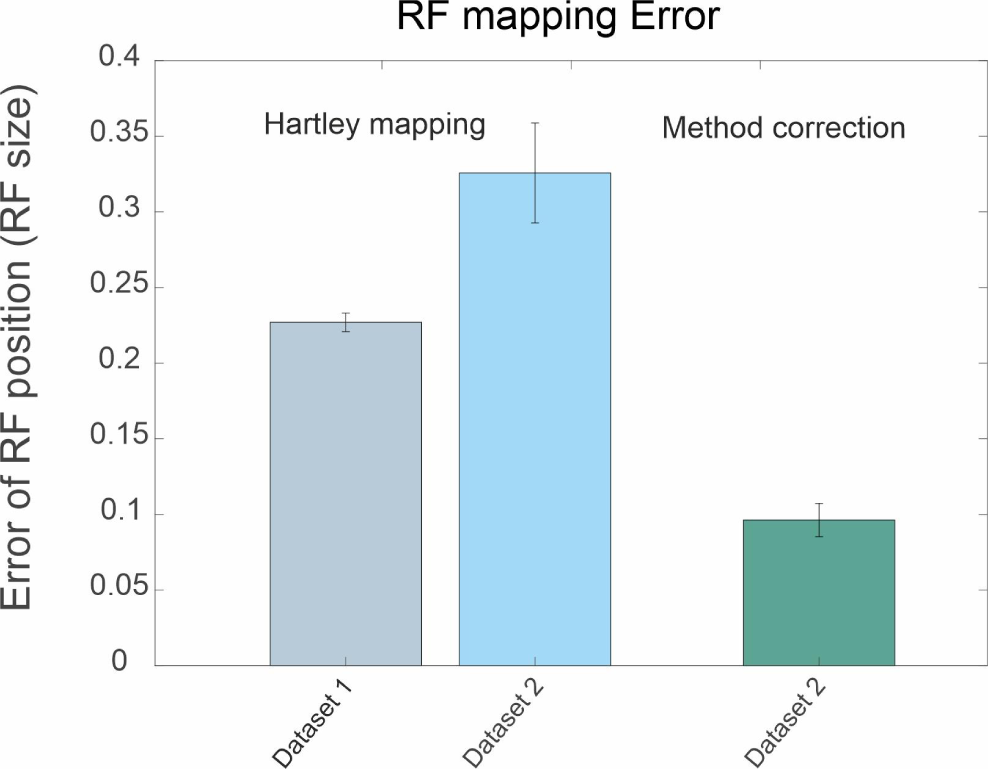
Error comparison between the geometric mapping and Hartley mapping. Gray column illustrates the average Hartley RF mapping error for 270 NOS monkey neurons, obtained by placing the drifting grating stimulus center at the center of the Hartley map. The spatial error is calculated based on the amplitude of sinusoidal patterns in the raster plot and normalized by RF size for each neuron. The blue column shows the average Hartley RF mapping error for 11 NOS Cat neurons, obtained by placing the drifting grating stimulus center at the center of the Hartley map. The green column is the average RF mapping error based on applying our method in the raster plot of 11 neurons of cats. The loads on each row represent the standard error of the mean (SEM). Using our method could reduce the mapping error by an average of one-third.

The reason behind the accuracy of our method relative to Hartley lies in the ability of our method to ignore the noisy nature of the neurons, and only attend to the overall patterns. On the other hand, Hartley takes into account every spike, which leads to more error when there is high spontaneous and baseline activity in the neuron. Moreover, Hartley might produce very misleading receptive fields if the used sorting algorithm fails in isolating neurons completely. However, using the fact that each NOS neuron produces sinusoidal raster plot patterns, we can detect such spike sorting failures if two sinusoidal patterns are detected in a raster plot, and further improve spike sorting as well as RF mapping. As another advantage of our method, we can find mapping errors based on sinusoidal patterns but reverse correlation mappings like Hartley only provide RF color maps and do not report any information for spatial error.

## Discussion

Response of V1 neurons to the orientation task (drifting grating ^18^ or moving bar ^17^) creates specific patterns in raster plots. Based on geometrical modeling and consistency of our model’s output with the real neuronal response, we have demonstrated that the key point behind these patterns is the relative location of the stimulus rotation center to the RF sub-regions.

Based on these patterns and our geometrical model, we proposed an accurate RF mapping method with an error of almost 1/3 of Hartley mapping. Previous RF mapping methods only result in a 2D RF map and do not provide any information about the spatial accuracy of obtained RF. But, based on the amplitude of sinusoidal patterns, our method can inform us about the spatial accuracy of obtained RF. For ease of implementation of this method, we have written practical applications in the MATLAB and GeoGebra platforms. Although here we have not shown the application of our method in RF mapping for complex cells, its operation domain is not restricted to simple cells and could be applied for complex cells too. For this purpose, it is necessary to use orientation tasks with moving bars ^17^ or drifting gratings with low spatial frequencies to create regular patterns in response raster-plot.

In non-orientation selective neurons, we have developed our geometrical model based on sensitivity to only one point (one circular region) on the screen. For orientation-selective neurons, our model operates based on sensitivity to two points (circular regions) on the screen. So, we can predict that neurons with sharp orientation tuning have two strong (equally balanced) sub-regions and in neurons with broad tuning curves, one region is dominant over the others. Our prediction is in agreement with reported results by Ringach ^26^ in monkey V1.

This is a widespread belief that orientation-selective V1 neurons have RFs with two or more ellipsoid (elongated) sub-regions, which the shape of these subregions brings the ability of orientation selectivity and the preferred orientation could be predicted from RF structure ^20–25^. This belief comes from the proposed model by Hubel & Wiesel ^27^ and states that LGN neurons with aligned center-surround RFs converge to an orientation-selective V1 neuron and is supported with later computational ^28–30^ and experimental studies ^17,31^. Also, most RF mapping tasks like Hartley lead to RF maps with elongated sub-regions for orientation-selective neurons^13,26,32^.

Based on this classical model ^27^, the existence of orientation-selective V1 neurons with only one elongated sub-region is possible. But such forms of RFs have not been observed yet and our model clearly explains why there is no orientation-selective V1 neuron with only one elongated sub-region?

Our geometrical model reveals that orientation selectivity in V1 neurons occurs based on two spatially offset opposing circular sub-regions and the real shapes of RF sub-regions are not elongated. This finding is congruent with previous reports by Jin and Lien ^25,31^ and also illustrates how orientation selectivity works in V1.

The ellipsoid (elongated) shape of RF sub-regions is because of the RF mapping task and not the real shape of sub-regions. For example, in orientation-selective V1 neurons, RF mapping with sparse noise results in circular sub-regions instead of elongated sub-regions ^13,17,25^. Based on the geometrical model, we can explain that this phenomenon occurs because of different stimulus ensembles used in RF mapping tasks. Indeed, RF mapping tasks in which the stimulus covers both sub-regions simultaneously result in elongated sub-regions (like Hartley stimulus ^9,13,26,32^). But, RF mapping tasks in which the stimulus covers only one sub-region at each time (each frame of the monitor) result in circular sub-regions (like sparse noise ^13,17,25^).

Ringach ^30^ explains the development of pinwheel orientation maps in V1, based on the arrangement of ON and OFF-center retinal ganglion cell mosaics. In this manner, our model can predict the existence of regularly arranged ON and OFF-center V1 cell mosaics in the cortex.

In the end, our model can perfectly reproduce the response of orientation-selective V1 neurons only based on sensitivity to two circular regions on the screen. So, based on this model we can theoretically propose that only 2 LGN neurons are needed for orientation selectivity in one V1 neuron. But, in the Hubel & Wiesel model, many LGN neurons are needed for orientation selectivity in one V1 neuron. Optimum usage of neurons is another advantage of our proposed model and can help us in better understanding LGN to V1 connections.

## Methods

In this paper, we used Adult male cats (ages: 6–10 months, n =6) and monkeys (Macaca mulatta, n = 2 males) V1 neuronal data. We have recorded 139 isolated cat and monkey neurons, and as shown here, there is a great agreement between the model and all of the neuronal responses. In another evaluation, we have shown (Supplementary) the perfect implementation of our method to find accurate RF locations in multichannel recordings (30 cat neurons previously recorded in Dr. Jose Manuel Alonso’s lab). Also, we used data from 270 Monkey neurons (previously recorded in Dr. Alonso’s lab) to calculate Hartley RF mapping error and evaluate our method. All procedures were performed in accordance with the guidelines of the Research Ethics Committee at the Institute for Research in Fundamental Sciences (IPM) and the U.S. Department of Agriculture and approved by the Institutional Animal Care and Use Committee at the State University of New York, State College of Optometry.

### Non-human primate experiments

Monkeys (Macaca mulatta, n=2 males; State University of New York, Optometry College) were implanted with a scleral eye coil to track eye movements, ahead post for head fixation, and a chronic array of electrodes was implanted in the primary visual cortex. Analgesics and antibiotics were administered before and after the surgery. The monkeys were trained to touch a bar and fixate on a small cross (0.1°/side) presented at a distance of 57 cm. They maintained the eye position within 0.5–1° for 3 s and released the bar when the cross changed color to receive a drop of juice as a reward. Details of visual stimuli and computer screens for Monkey’s experiments have been described previously [15, 16].

### Anesthetized cat experiments

Acepromazine (0.2 mg/kg) and Atropine (0.05 mg/kg) were injected initially for relaxation and sedation of the animal and decrease of saliva secretion. Initial anesthetization was begun with IM injection of Ketamine (20mg/kg). After shaving both legs of the cat, cannulations of both femoral arteries were performed for delivering drugs. Dexamethasone (0.15 mg/kg) was injected (IM) in order to reduce inflammation. Dexamethasone and atropine were injected every 8 hours. An endotracheal intubation was performed to connect to the mechanical ventilator. Lidocaine HCL (2%) was applied to all incisions and pressure points of the stereotaxic in contact with the animal’s body. Cat’s head was supported non-traumatically in the stereotaxic. General anesthesia was induced with intravenous Sodium thiopental (3-8 mg/kg/h) solution (diluted with normal saline) and maintained with a continuous infusion of thiopental. All vital signs were closely monitored and carefully maintained within normal physiological limits. Before starting craniotomy, fentanyl (3mcg/kg/h) was injected to decrease surgical pain. The skull and dura (10 mm D) overlying the primary visual cortex were removed. After the surgery, Cis-atracurium (0.18mg/kg) was injected initially and the animal was connected to a ventilator with a respiration rate of 28 breaths per minute performed. Paralysis was maintained during recording using a continuous infusion of Cis-atracurium (0.54mg/kg/h). A potting eye-opener, then nictitating membranes were retracted with 2% neo synephrine and the pupils dilated with 10% atropine sulfate, and the eyes were fitted with gas-permeable contact lenses (+2.0 D) [17] to focus on a tangent screen. By using a fiber-optic light source, the positions of the optic disk and the area centralis were plotted on the tangent screen that was placed 57 cm in front of the animal. The weight of the animals ranged from 3.2 to 4.4 kg and the ages ranged from 6 to 10 months (n=6). After lowering the electrodes to the cortical surface, agar is used to insulate the cortex.

### Electrophysiological recordings and data acquisition

To measure the activity of single isolated V1 neurons, the single electrode was slowly lowered inside the primary visual cortex while the Hartley stimulus, the static grating stimuli with a wide range of properties (spatial frequency, orientation and phase) was presented in a CRT monitor. We subjectively detect the neurons from their clear firing rate increment as a consequence of stimulus presentation; the firing rate increase was audible through the speakers. We then isolated the neurons manually, using Blackrock software. The neuronal activity was recorded using the FHC tungsten single-electrode and the Black-rock data acquisition. The results were filtered online and in the frequency range of 500 to 6,000 Hz, the spikes are separated using a manual threshold. The neuronal spiking activity was displayed on a computer for online analysis. Online display of results was essential for designing exclusive stimuli for each isolated neuron. Also, the results were obtained after offline filtering and the accurate spike sorting have been obtained semi-manually using neuronal networks classifiers. All the analysis was performed under MATLAB software and the codes used for the analysis belong to the laboratory.

### Mapping visual response properties of V1 neurons

The Visual stimuli were displayed on the Hansol 920D monitor with, 75Hz refresh rate, resolution of 1280 × 960 pixels, Luminance of 56 Cd/m2, and dimensions of 27 36 Cm, which was 57Cm away from the subjects’ eyes. Using Psychtoolbox, we designed the optimum specific stimulus for each neuron to perfectly activate it. Also, the stimulus presentation timing was checked frame by frame using a photodiode, with timing error less than 0.0001 seconds in one second of the presented stimulus. At the beginning of the experiment, we used drifting grating stimulus manually to estimate the approximate location of the receptive field and the initial features of the single isolated neuron (Approximate Receptive-Field size, Spatial Frequency, and Orientation). According to the initial finding of response parameters, we generated a set of full-field gratings to obtain the appropriate response activity (e.g., 18 orientations, 8 spatial phases, and 6 spatial frequencies (0.03: 0.3 cycles/deg). Then we applied features to generate Hartley stimuli to determine the neuron’s receptive field and obtain optimum response properties. Images of Hartley stimulus were displayed in 960 by 1200-pixel quality and the display frequency is half the monitor frequency with 75 images in 2 seconds (37.5 Hz). By placing the optimum drifting gritting stimulus at the center of the Hartley map, we plot neurons’ response as the spike raster plot. We also performed our calculations based on the monitor pixels so that the results could be easily used to modify the placement of the stimulus. To convert the dimensions to the degree of vision, according to the distance of the monitor, every 1 cm of the monitor is equal to 1 degree and 960 pixels is equal to 27 cm.

### Experimental design (orientation task)

The orientation task was presented as moving bars ^17^ or a drifting gritting stimulus with the optimum spatial frequency (obtained based on Hartley results) and drifted at the speed of 3 cycles per second, which was displayed at angles of 0 to 360 with 10 degrees for each step. Drifting grating with the speed of 3 cycles per second in contrast to the speed of 2 c/s reduces the neuronal response variability and therefore, by increasing the quality of patterns, would increase the accuracy of the method. Drifting gratings were presented for 4 seconds and three repetitions per angle (0:10:360 degree fig. 1a). In total, each spike raster plot consists of 108 trials. We also determined the size of the stimulus according to the result obtained from the Hartley task to have the maximum firing rates. To fasten the orientation task, reduce the duration of each trial to less than 1 second in proportion to the drift speed. Also, it is not necessary to repeat each trial 3 times. We used these parameters only to increase the accuracy of the results. In fact, we can obtain sinusoidal spike raster patterns without repetition of trials and at the shorter time of stimulus presentation. In order to use our method in multi-electrode recordings and mapping multiple neurons simultaneously, we propose using the orientation task with moving bars^17^.

### Reverse correlation Mapping

The drifting grating task is made up of a set of grating images that are displayed frame by frame on the monitor with a slight incremental phase difference relative to each other. In fact, the drift phenomenon in the drifting grating task is caused by the arrangement of several grating images in consecutive frames that the phase of these images increases frame by frame in time. Due to the high frame rate (75 Hz), we see a continuous movement (drifting). Therefore, we have obtained accurate receptive field maps, using drifting grating images and their respective neuronal response in reverse correlation. In these maps, we used a color map with blue and red colors (fig. 4a/c middle). Blue and red colors indicate the sensitivity of the neuron to black and white stimulus on the screen.

### Driving of geometrical equations

This table represents the geometrical descriptions of the suggested model to calculate the cell’s response latency relative to the rotation angle (fig. 2d).

## Supplementary information

**Figure 1:**
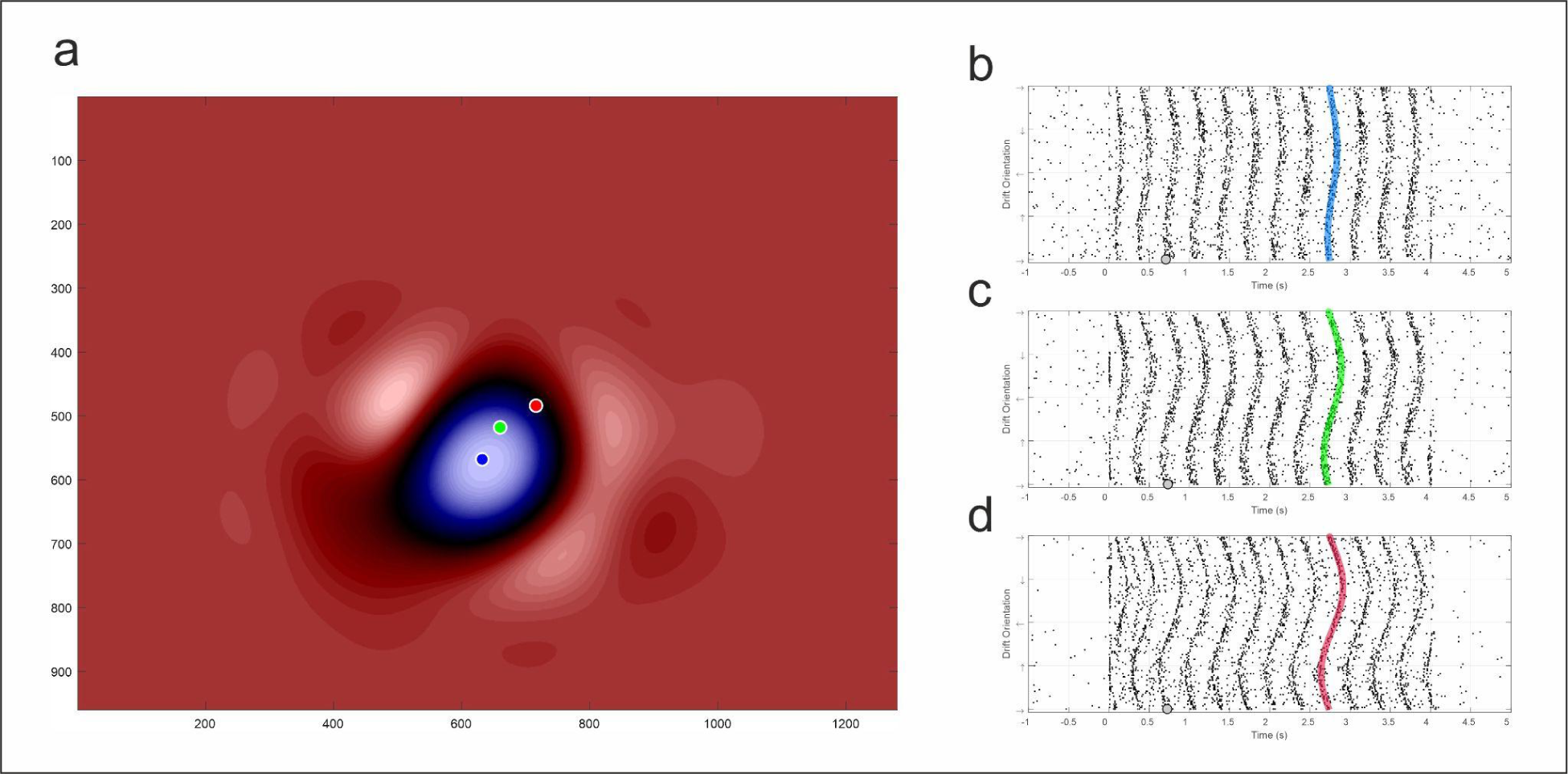
Increasing the distance between the stimulus rotation center to the RF center increases the amplitude of sinusoidal patterns in the raster plot. (a) RF map for a NOS neuron (OFF center) obtained by Hartley mapping. Blue, green, and red points indicate the location of the drifting grating rotation center in b, c, and d records. As shown, by distance from the RF center, the amplitude of sinusoidal patterns increases.

**Figure 2:**
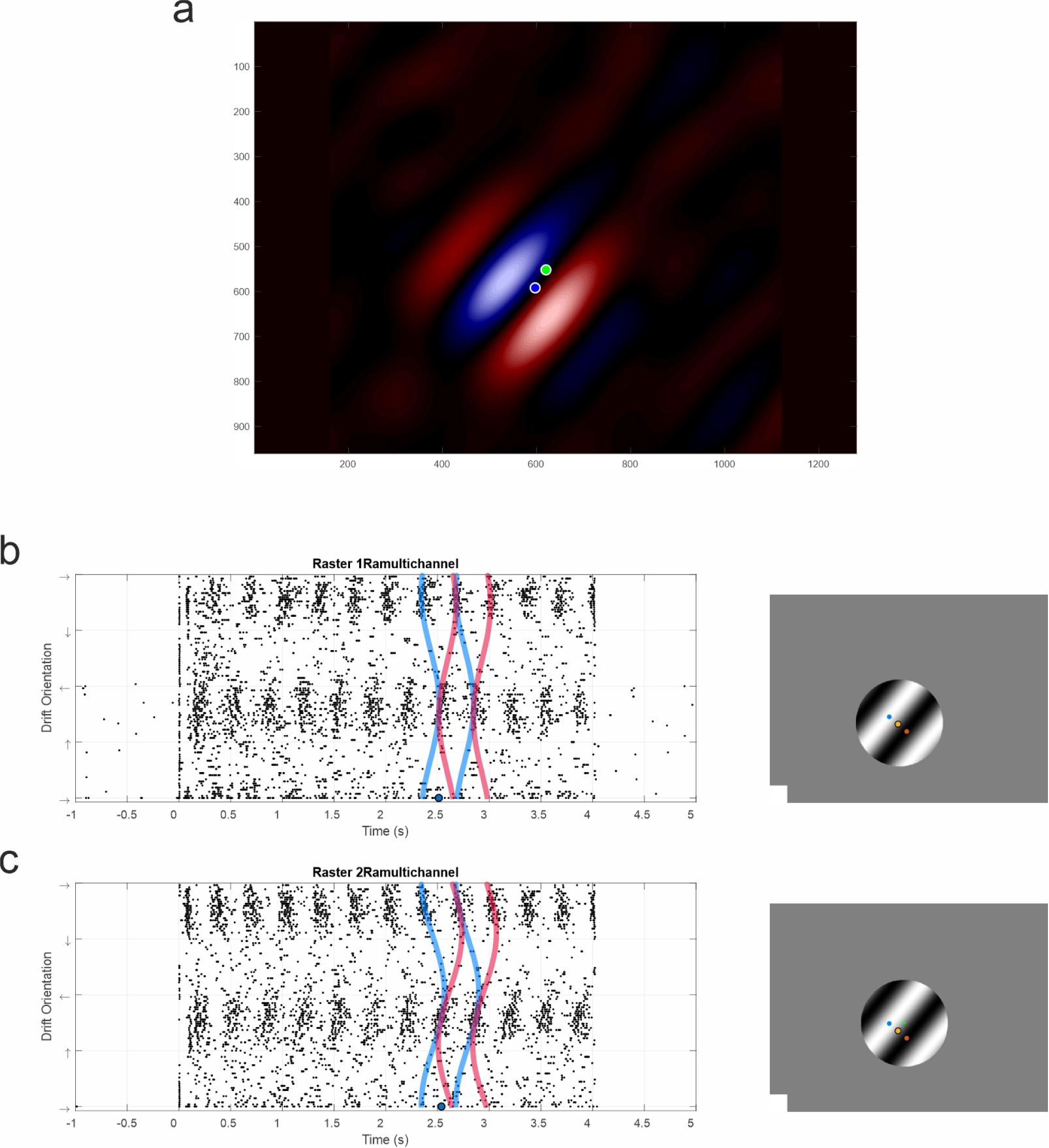
Different stimulus rotation center positioning creates different patterns in the raster plot. (a) RF map of an OS neuron obtained by Hartley mapping. (b) Positioning the grating rotation center (green point, right) at the center of RF (blue point in (a) and yellow point in (b), right) create patterns as predicted by the model (fig.5.b). (c) Distancing upwards from the RF center (green point in (a)) creates oblique patterns in the raster plot. The shape of these patterns depends on the relative positioning of the stimulus rotation center and the RF center. As shown, by fitting the model’s output on the raster plot patterns, we can find the exact location of RF sub-regions accurately.

**Figure 3:**
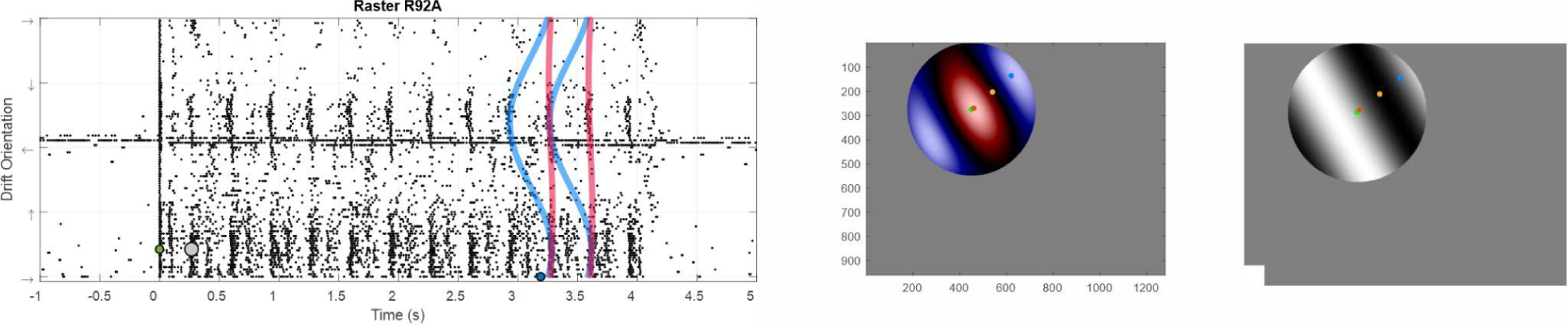
positioning of stimulus rotation center on the RF sub-region. Positioning the grating rotation center (green point right) on the center of one of the sub-regions creates such vertical patterns in the raster plot. The output of the model fitted on the pattern and also, location of RF sub-regions (blue and red dots, middle and right) are completely matched to the reverse correlation mapping method.

